# Balancing selection drives maintenance of genetic variation in *Drosophila* antimicrobial peptides

**DOI:** 10.1101/298893

**Authors:** Joanne R. Chapman, Tom Hill, Robert L. Unckless

**Author notes:** These authors contributed equally to the work (Joanne R. Chapman, Tom Hill). The *Drosophila* sequence data were obtained from John Pool’s *Drosophila* Genome Nexus http://www.johnpool.net/genomes.html).

## Abstract

Genes involved in immune defense against pathogens provide some of the most well-known examples of both directional and balancing selection. Antimicrobial peptides (AMPs) are innate immune effector genes, playing a key role in pathogen clearance in many species, including *Drosophila*. Conflicting lines of evidence have suggested AMPs may be under directional, balancing or purifying selection. Here, we use a case-control gene approach to show that balancing selection is an important force shaping AMP diversity in two species of *Drosophila*. In *D. melanogaster*, this is most clearly observed in ancestral African populations. Furthermore, the signature of balancing selection is even clearer once background selection has been accounted for. Balancing selection also acts on AMPs in *D. mauritiana*, an isolated island endemic separated from *D. melanogaster* by about 4 million years of evolution. This suggests that balancing selection may be acting to maintain adaptive diversity in AMPs in insects as it does in other taxa.

## 1 Introduction

Pathogens exert strong selective pressures on their hosts, both in terms of individual fitness and the evolutionary trajectory of populations and species. Co-evolutionary dynamics of hosts and pathogens results in continual selection for adaptive improvements in both players, often referred to as a co-evolutionary arms race (1, 2, 3). As a consequence, genes involved in immune defense tend to undergo strong positive selection, such that they are among the fastest evolving genes in the genomes of many hosts (4, 5, 6, 7, 8).

However, resistance mutations may not always become fixed. Balancing selection is the process whereby polymorphism is adaptively maintained within genes over extended timescales, sometimes described as trench-warfare dynamics (9). Several processes are thought to contribute to balancing selection (reviewed in (10)). These include heterozygote advantage, whereby individuals heterozygous at a given locus have a fitness advantage over either homozygote; negative frequency dependent selection, whereby the benefit of an allele increases the rarer it is in a population; and selection varying in a context-dependent manner, for example at different spatial or temporal scales, between the sexes, or in the presence or absence of infection. Balancing selection can be detected as an excess of intermediate frequency variants and a region of increased polymorphism around the selected site. The extent to which selection will impact genetic variation within and around immune genes will depend on a number of factors, including: the timescale upon which selection is acting (11); the density, diversity and virulence of pathogens (12); the cost of maintaining resistance alleles in the absence of infection (13); effective population size, mutation and recombination rates of hosts and pathogens (14); environmental variables (15); and demographic factors such as gene flow and bottlenecks (16).

The dynamic selective pressures exerted by pathogens promote balanced polymorphism of host immune genes in several cases. Perhaps the best documented example is the major histocompatibility complex (MHC) in vertebrates (reviewed in (17, 18, 19, 20)). Individuals tend to be heterozygous at MHC loci, and large numbers of MHC alleles are maintained in populations. Other examples of balancing selection acting on host immune genes in animals include toll-like receptors (TLRs) in humans (21), red deer (22) and birds (23, 24); various cytokine genes (particularly interleukins) in humans (21, 25, 26, 27), birds (28, 29, 30) and voles (31); and viral resistance genes including *Oas1b* in mice (32), *OAS1* in primates (33, 34) and *TRIM5* in humans (35) and primates (36). Balancing selection also appears to play a role in the evolution of antimicrobial peptides (AMPs). AMPs are effectors of innate immunity that are strongly induced upon infection (37, 38). They tend to be membrane active (39, 40), with a direct role in killing and/or impeding the growth of pathogens (41, 42). Balancing selection has been implicated as a driver of AMP evolution in a diverse array of species including birds (43, 44), amphibians (45), fish (46), molluscs (47) and humans (48, 49).

The fruit fly, *Drosophila melanogaster*, is an important model for understanding evolution of the immune system (50, 51, 52, 53, 54). Directional selection on *Drosophila* immune genes appears to be a relatively widespread phenomenon, especially amongst receptor and signaling genes (55, 56, 57, 58, 59, 60). In contrast, evidence for balancing selection acting on *Drosophila* immune genes has been more equivocal. Genome-wide scans by Croze and colleagues (61, 62) found little evidence for balancing selection acting on immune genes in general, and Obbard *et al.* (58) found no evidence for adaptive evolution of AMPs. In contrast, both single gene and genome-wide analyses of selection have indicated that balancing selection (13, 63) or diversifying selection (64) may play an important role in the evolution of AMPs in *Drosophila*. Additionally, recent analyses have shown that seasonal fluctuations in temperate can cause rapid oscillations in *D. melanogaster* allele frequencies (65), particularly in immune genes, including AMPs (66, 67).

Insects and other invertebrates lack an adaptive immune system, so AMPs play a key role in controlling pathogen load and infection outcome (41, 42). Given their direct interaction with pathogens, it is surprising that AMPs do not show signatures of recurrent adaptive substitutions. We hypothesize that AMPs in insects are prone to balancing selection. To test this hypothesis, we examined AMP variation in four populations of *Drosophila melanogaster* and one population of *Drosophila mauritiana*. Using a case-control gene approach, we searched for molecular evolutionary signatures of selection. Our results provide evidence that balancing selection is an important driver of AMP evolution.

## 2 Results

### 2.1 Genetic variation across four *Drosophila melanogaster* populations

To determine whether AMPs show signatures of balancing selection, we examined nucleotide polymorphism data in *D. melanogaster* populations. Coding sequence alignments for 13494 genes (including 35 AMPs and 104 immunity genes) were obtained (68) for four *D. melanogaster* populations: Zambia (ZI), Rwanda (RG), France (FR), and North Carolina (DGRP) (see Materials and Methods, Supplementary Table 1). *D. melanogaster* originated in Sub-Saharan Africa, expanded into Europe approximately 15-16,000 years ago, and subsequently spread to North America less than 200 years ago (69, 70, 71). The ZI and RG lines therefore represent ancestral populations, whereas FR and DGRP are derived populations.

We calculated three population genetic statistics: Watterson’s *θ* (the sample size corrected number of segregating sites), *π* (pairwise nucleotide diversity) and Tajima’s D. Consistent with balancing selection occurring in AMPs, the mean Tajima’s D for AMPs is higher than the average across autosomes for Zambia (ZI, −0.713 AMPs versus −1.168 autosome average), Rwanda (RG, −0.358 versus −0.503), France (FR, 0.033 versus −0.021), and the DGRP (−0.171 versus −0.179, Supplementary Table 2). As observed previously (e.g. (72, 73)), the autosome-wide average for Tajima’s D is quite negative in *D. melanogaster*, which likely reflects a complex demographic history. In general, a significantly higher proportion of AMPs have a positive Tajima’s D when compared to other genes on autosomes (Supplementary Table 3; *χ*^2^ *p*-value < 0.02 for all populations except France where *χ*^2^ *p*-value = 0.36).

### 2.2 Case-control tests for balancing selection in *Drosophila*

Given the apparent differences in selection between AMPs and the genome averages described above, we employed a case-control approach to test whether AMPs showed signatures of balancing selection while controlling for local variation in mutation and recombination rates. For each AMP, we randomly sampled genes of similar length (amino acid sequence length *≤*10 times the size of the AMP) and position (within 100kb on either side), calculated statistics for the AMP and control gene, and then calculated the mean difference over the 35 AMP/control comparisons. We repeated this 10000 times to obtain an empirical distribution of differences (Figure 1). In these instances, a positive difference suggests a higher value for AMPs versus the control gene, and therefore a role for balancing selection. These differences are primarily positive for both *π* and Watterson’s *θ* for all populations (Figure 1B-C, Table 1). For Tajima’s D, the differences are positive for Zambia and Rwanda (ancestral populations), supporting balancing selection, but close to zero for France and negative for the DGRP (derived populations, Figure 1A, Table 1).

**Figure 1:**
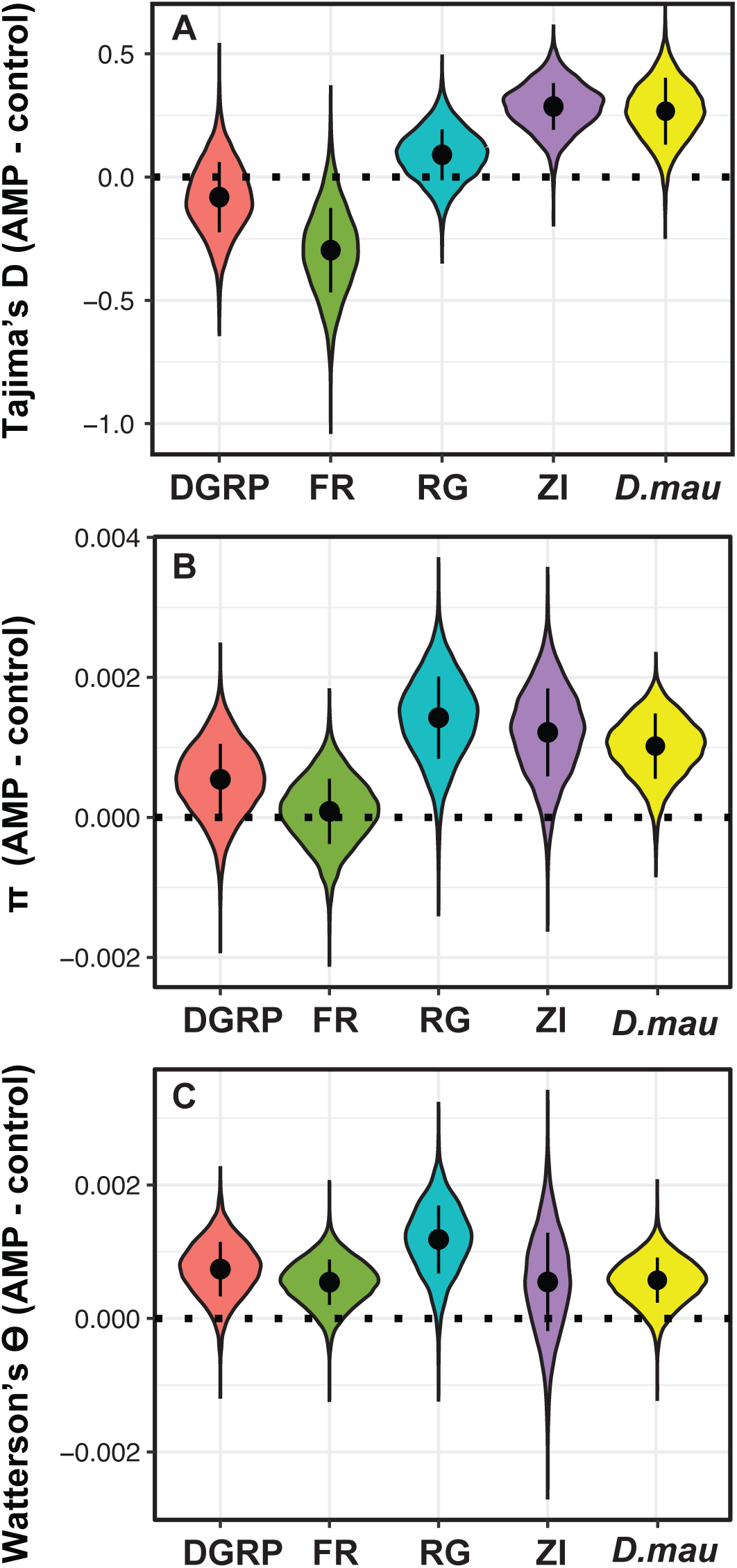
Difference in means between 35 AMPs and randomly chosen control genes, resampled 10000 times, separated by population (DGRP = Drosophila Genetics Reference panel from North Carolina, USA; FR = France; RG = Rwanda; ZI = Zambia). A) Tajima’s D, B) *π*, C) Watterson’s *θ*. The black dot within each plot shows the median for that population, and the black bar around the dot visualizes the interquartile range of the distribution.

**Table 1:** AMP minus control gene differences for three statistical measures of selection in four *D. melanogaster* populations and one *D. mauritiana* population. First row per statistic: percentage (%) of 10000 replicates in which the AMP minus control difference was positive (>0), suggestive of balancing selection; second row: mean AMP minus control difference across 10000 replicates; third row: standard deviation (std. dev.) of the mean (DGRP = Drosophila Genetics Reference Panel from North Carolina, USA; FR = France; RG = Rwanda; ZI = Zambia).

| AMP - control statistics | DGRP | FR | RG | ZI | *D. mauritiana* |
| --- | --- | --- | --- | --- | --- |
| Tajima's D differences $> 0$ (%) | 28.7 | 4.1 | 81.4 | 99.7 | 98.1 |
| Tajima's D differences mean | -0.084 | -0.295 | 0.092 | 0.289 | 0.26 |
| Tajima's D differences std. dev. | 0.142 | 0.171 | 0.102 | 0.092 | 0.12 |
| $\pi$ differences $> 0$ (%) | 85.9 | 58.4 | 98.9 | 96.9 | 100 |
| $\pi$ differences mean | 9.6e-5 | 9.6e-5 | 1.4e-3 | 1.2e-3 | 1.2e-5 |
| $\pi$ differences std. dev. | 5.5e-4 | 4.8e-5 | 5.5e-3 | 6.1e-4 | 2.1e-6 |
| Watterson's $\theta$ differences $> 0$ (%) | 96.2 | 93.7 | 98.5 | 77.4 | 99.9 |
| Watterson's $\theta$ differences mean | 7.5e-4 | 5.5e-4 | 1.2e-3 | 5.6e-4 | 1.7e-5 |
| Watterson's $\theta$ differences std. dev. | 4.1e-4 | 3.4e-4 | 5.1e-4 | 7.4e-4 | 1.5e-6 |

To identify if these signatures of balancing selection are unique to AMPs, or consistent across all immunity genes, we repeated all tests, this time for all non-AMP immunity genes. We found very little evidence of balancing or directional selection across the remaining immunity genes, with differences closer to zero (Supplementary Tables 2 and 4, Figure S1). This result is in general concordance with those of Croze *et al* (61, 62).

It is possible that the observed signature of balancing selection amongst AMPs is due to various sampling artifacts. First, AMP families tend to occur in clusters throughout the genome, so it is possible that including all AMPs in the analyses effectively counts the same selective event multiple times. To account for this, we subsampled 10 unlinked (>5kb apart) AMPs and repeated our analyses. This did not qualitatively change our results (Supplementary Figure 2). Second, the presence of the selfish genetic element *Segregation Distorter* (*SD*), a low-frequency autosomal meiotic drive element (74) on the second chromosome, in some lines (4% in both Zambia and France) may influence our results. However, removing these lines did not qualitatively change our results (Supplementary Table 5, Supplementary Figure 3). We therefore consider that the observed patterns reflect true underlying evolutionary processes rather than sampling artifacts.

### 2.3 Accounting for background selection strengthens the signature of balancing selection on *Drosophila* AMPs

Background selection, the removal neutral variation due to selection against linked deleterious alleles, can influence levels of polymorphism across the genome. Comeron (75) calculated the observed amount of background selection across the genome in 1000 base pair (bp) windows in the Rwanda population. He then correlated silent polymorphism against this measure. Regions with positive residuals (more silent polymorphism than expected based on background selection) were deemed to be under balancing selection, while those with negative residuals (less silent polymorphism than expected based on background selection) were deemed to be under directional selection. Two regions that contain AMPs (IM4 and Cecropin) were among the handful of outliers discussed by Comeron as being under balancing selection. We identified all AMP-containing windows and replotted Comeron’s data. This revealed that AMPs tend to fall in regions well above the trend-line (red points, Figure 2A), indicating they are evolving in a manner consistent with balancing selection. To further ascertain whether AMPs as a group show signatures of balancing selection, we used Comeron’s background selection data (75) to calculate residuals for regions containing AMPs and compared them to residuals for randomly chosen position- and size-controlled genes employing methods similar to those used in the previous analyses. The distribution of differences in residuals was always above zero (Figure 2B, mean = 1.63, std. dev. = 0.20). This supports Comeron’s assertion that accounting for background selection improves the ability to detect balancing selection, and also supports our previous results showing that AMPs as a group are subject to balancing selection.

**Figure 2:**
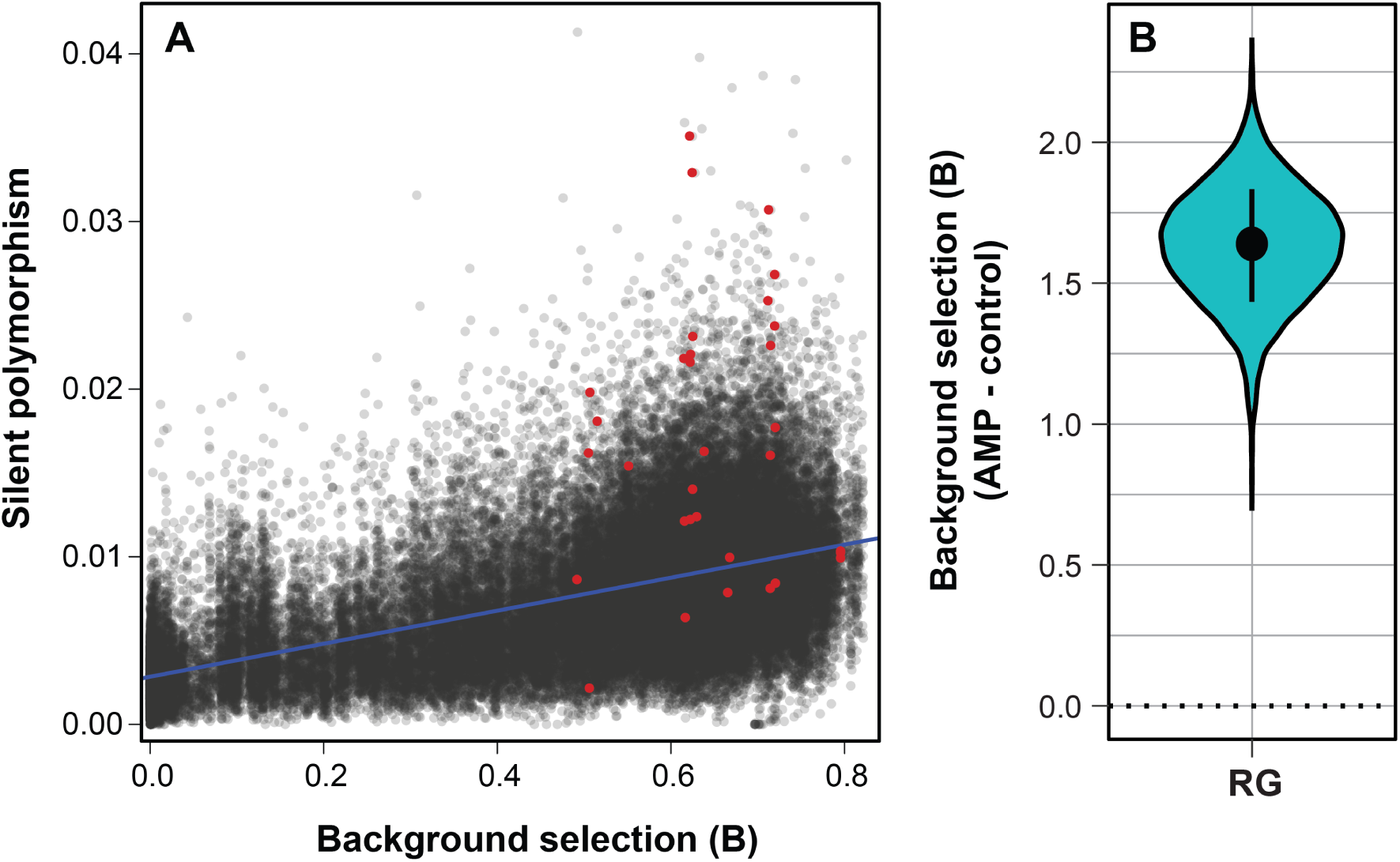
Accounting for background selection strengthens the signal of balancing selection on AMPs. A) Correlation between silent polymorphism and background selection (B) in 1000bp windows for the Rwanda population of *D. melanogaster*. The line of best fit is in blue and regions containing AMPs are indicated by red dots, B) Resampling of mean difference in the background selection statistic between AMPs and control genes.

### 2.4 Balancing selection also acts on *Drosophila mauritiana* AMPs

We also calculated population genetic statistics for 9980 genes in 107 *D. mauritiana* isofemale lines, sequenced as a pool. *D. mauritiana* is an island endemic which diverged from *D. melanogaster* approximately 3-5 million years ago (76, 77). SNP frequencies were called using Popoolation which accounts for low frequency variants and variation in coverage that may influence results from pooled samples (78). As found for *D. melanogaster*, there was a significant excess of AMPs with a positive Tajima’s D compared to all other genes (*χ*^2^ = 19.96, *p*-value < 0.0001), and AMPs have a higher mean Tajima’s D (−1.034 versus −1.463). We again resampled the difference in these statistics between AMPs and neighboring control genes. We found AMPs have consistently higher values for *π*, Watterson’s *θ* and Tajima’s D than their matched controls (Figure 1, Table 1, Supplementary Tables 2 and 3, Supplementary Figure 4). For other immunity genes, the differences from controls are primarily negative for *π*, Watterson’s *θ* and Tajima’s D, suggesting directional selection may be acting on these genes (Supplementary Table 4, Supplementary Figure 4) in *D. mauritiana*.

## Discussion

We find evidence consistent with balancing selection being an important evolutionary driver of AMP genes in *Drosophila*. This is most clearly observed in ancestral African populations (Zambia and Rwanda). There are several reasons why previous analyses may not have conclusively identified the selective forces acting on AMPs. First, signals of selection can be clouded by background selection. We found that the clearest signal for AMP balancing selection was in the Rwandan population after using Comeron’s method (75) to account for background selection. Second, previous studies have tended to group immune genes as a single entity when scanning genomes for footprints of selection. Strong directional selection acting on some receptor and signaling immune genes may swamp a subtler signal of balancing selection acting on antimicrobial peptides. Third, this effect may be exacerbated by the fact that effector genes tend to be smaller (42) than receptor and signaling genes. Fourth, patterns of nucleotide polymorphism are strongly influenced by population demographic history. Our case-control approach should account for the confounding influences of local mutation and recombination rate variation, gene size and demography (79).

As populations establish in new habitats the pathogen pressure will be different, as will prevailing environmental conditions. This could dramatically alter which alleles are selectively advantageous. Loss of disadvantageous alleles (for example alleles resistant to pathogens not present in the new habitat) likely occurs more rapidly than establishment of new, beneficial polymorphisms (for example resistance alleles for newly encountered pathogens). This may explain why we find the strongest evidence for balancing selection on AMPs in ancestral African populations that have been co-evolving with their pathogens, under semi-predictable conditions, for long time-periods.

It is tempting to look to newly developed methods for detecting balancing selection (80, 81), but these statistics were developed for detecting the molecular footprints of selection in human populations. Assumptions about the genomic signatures of a balanced polymorphism that work well in humans are not applicable to *Drosophila*, because the window of linked polymorphism likely to show these signatures is tiny. To state this numerically, DeGiorgio *et al.* (81), based on Gao *et al.* (82), suggest a window size of 1/*ρ* (where *ρ* is the population-scaled recombination rate or 4*N*_*e*_*r*) for observing the signature of a linked balanced polymorphism. For humans, *ρ* is about 0.001 so the window size is about 1000 bp (81). Estimates of *ρ* in *Drosophila* are highest in the DGRP population and range from 9.6 to 14.8 for the different chromosomes (83). These values correspond to windows of less than 1/10 of a single base in *Drosophila*, rendering these tests unusable in this genus.

We find that, at least in ancestral populations, AMPs tend to evolve in a manner consistent with balancing selection. This is in contrast to other immune genes that show no such pattern. Why are AMPs different than other immune genes? One characteristic of AMPs is that they interact directly with microbes (84), and, in some cases, AMP sequence is directly linked to the efficacy of bacterial membrane interactions (85). If particular AMP alleles encode for peptides that are more effective at fighting infection by particular microbes, a fluctuating suite of pathogens in the environment over time or space could lead to balanced polymorphisms. This “specificity hypothesis” suggests that allele frequencies in AMPs should vary spatially or temporally. There is some evidence for both seasonal (66) and spatial (67) variation in selection pressure on AMPs. However, evidence for AMP specificity against particular pathogens, especially different naturally occurring alleles of the same AMP, is currently rare (but see e.g. (63, 86, 87, 88)).

Alternatively, AMP variation might be maintained because AMP alleles that are more effective against pathogens also carry a higher autoimmune cost. This “autoimmune hypothesis” states that more effective AMP alleles should be common during pathogen epidemics, but decrease in frequency when pathogens are rare. These patterns might also vary spatially and temporally, making the interpretation of these context-dependent patterns more difficult. There is evidence that overexpression of AMPs can have deleterious fitness consequences (89, 90, 91). However, it seems that if autoimmune costs were important in maintaining variation, we would also see signatures of balancing selection in the IMD and toll pathway signaling genes that control expression of AMPs. Most work suggests that these genes are evolving under the arms race model (57, 58, 59). Distinguishing between these two hypotheses for the adaptive maintenance of AMP genetic variation will take careful functional analysis.

## 4 Methods

### 4.1 Polymorphism in four populations of *Drosophila melanogaster*

We downloaded chromosome sequences for the Zambia (ZI, n=197), Rwanda (RG, n=27), *Drosophila melanogaster* Genetic Reference Panel (DGRP, n=205) and France (FR, n=96) populations, available as part of the *Drosophila Genome Nexus*, from http://www.johnpool.net/genomes.html (92, 93). We then converted these sequences into FASTA files, per chromosome, for each population. We also created a second set of FASTA files that excluded chromosomes known to contain the *Segregation Distorter* (*SD*) haplotype (taken from: (74)). The RG and ZI populations are much higher quality data, the average per base coverage of the raw FASTQ data used to generate the FASTA files is much higher, and the number of ambiguous bases is much lower than the DGRP and FR populations (Supplementary Table 1).

Using annotation 5.57 of the *D. melanogaster* genome, we extracted the FASTA alignments for each gene. Following this, we used a custom bioperl script, with the the package Bio, to find *π*, Watterson’s *θ*, Tajima’s D and the number of segregating sites for each gene. We categorized each gene using the designations found in Obbard *et al.* (57). We removed non-autosomal genes from all downstream analyses, because the X chromosome does not harbor any AMPs.

For each analysis, (per population, including and excluding SD chromosomes) we then resampled to find the average difference in scores between case and control genes. Case genes were either a) AMPS, or b) immunity genes (using gene ontologies previously described (58)). For each gene in these categories, we randomly sampled a control gene within 100kbp upstream or downstream, that was no more than ten times larger than this gene and not another gene in the given category (AMP or immunity). We then found the average difference 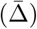 in each measure for the case (AMP/immunity) group and the control group such that:

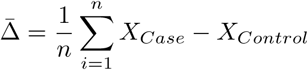

where *X*_*Case*_ represents the chosen gene, *X*_*Control*_ represents the randomly sampled control gene and *n* accounts for the number of genes in the group. We then repeated this 10000 times to obtain an empirical distribution of the differences.

We employ this method to control for genomewide variation in recombination rates, mutation rates, and possibly, demographic history. Resampling 10000 times allows for a robust empirical distribution that does not rely on the particular control genes chosen. We therefore present the distribution of differences as violin plots and purposefully do not discuss significance in terms of *P*-values. Instead, the proportion of resamplings that do not overlap zero is more analogous to a bootstrap value.

### 4.2 Polymorphism in a population of *Drosophila mauritiana*

We downloaded the reference genome, annotation and mapped BAM file of a population of *D. mauritiana* from http://www.popoolation.at/mauritiana_genome/, and used Popoolation to calculate Tajima’s D, *π* and Watterson’s *θ* for each gene in this population. We then resampled to find the average difference in scores between AMPs and a control set of genes, as described above.

## 5 Supplementary Material

**Supplementary Table 1** - Summary statistics for each dataset, including the average base coverage for each population and the average number of ambiguous bases per 1000 bases in the FASTA files used. Data taken from johnpool.net/genomes.html.

**Supplementary Table 2** - Summary statistics for each AMP and immunity gene for each population, also the mean for each statistic for all non-AMP immune genes.

**Supplementary Table 3** - *χ*^2^ test contingency tables for each population, showing the number of AMPs and other genes with positive and negative Tajima’s D.

**Supplementary Table 4** - Summary of resampling results across case (AMPs/immunity) genes and their matched control genes. These statistics include the percentage greater than 0, mean and standard deviation for each resampling set.

**Supplementary Table 5** - Summary of resampling results across case (AMPs/immunity) genes and their matched control genes, with all SD containing samples removed. These statistics include the percentage greater than 0, mean and standard deviation for each resampling set.

**Supplementary Figure 1.**
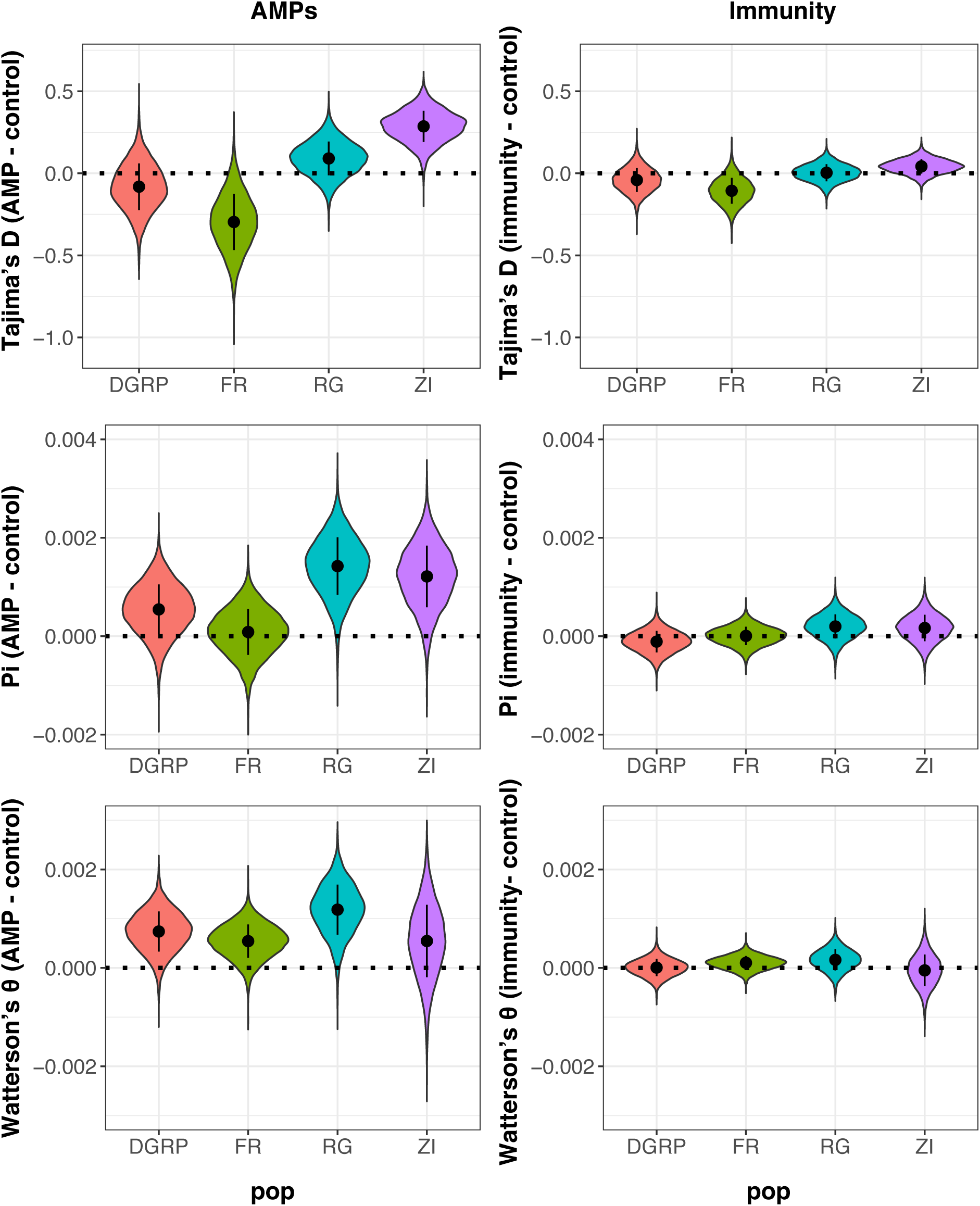
Summary of resampling results (case - control) for AMPs and other immunity genes for Tajima’s D, *π*, Watterson’s *θ*

**Supplementary Figure 2.**
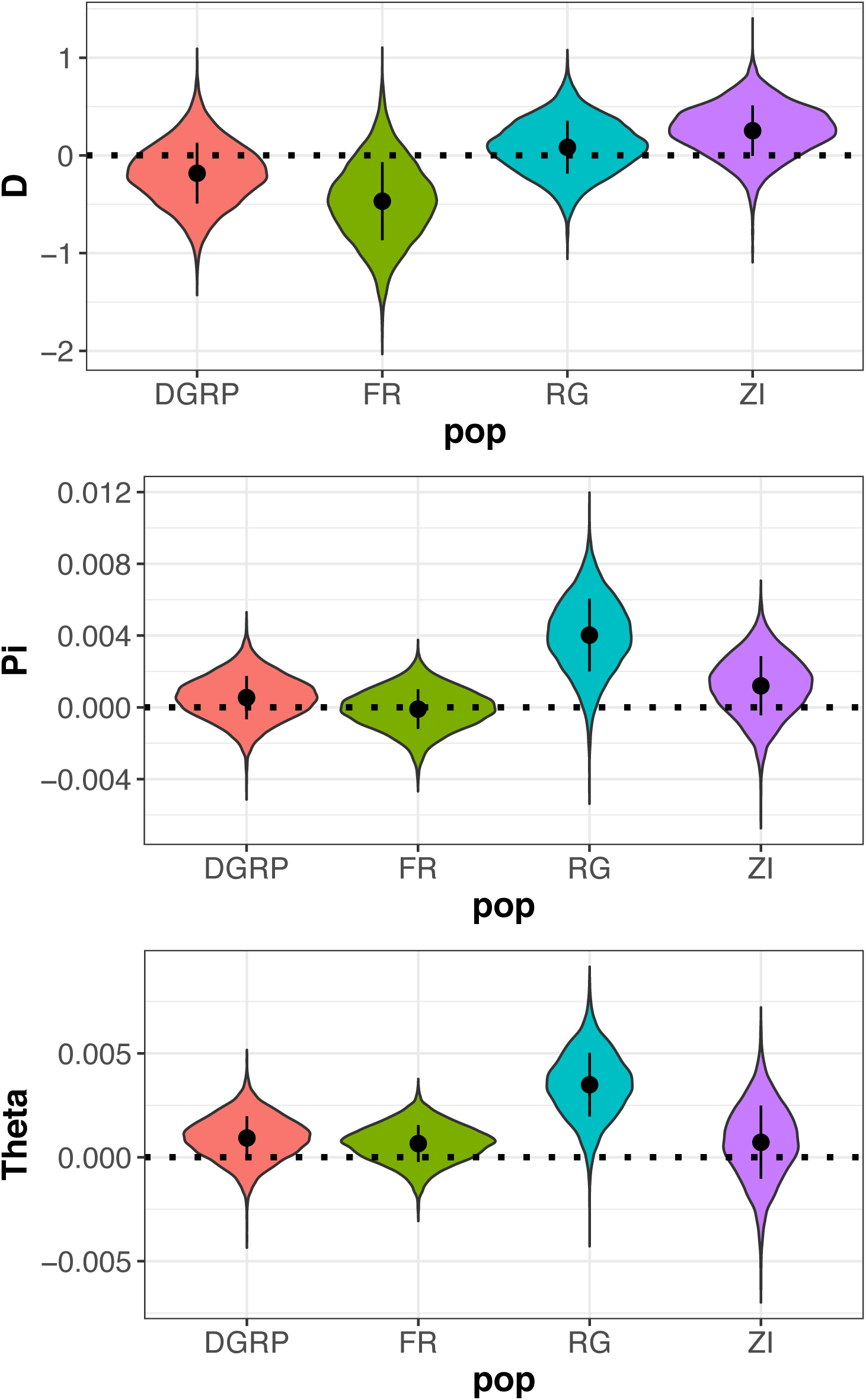
Summary of resampling results (case - control) for AMPs for Tajima’s D, *π*, Watterson’s *θ*, using the subset of non-linked AMPs.

**Supplementary Figure 3.**
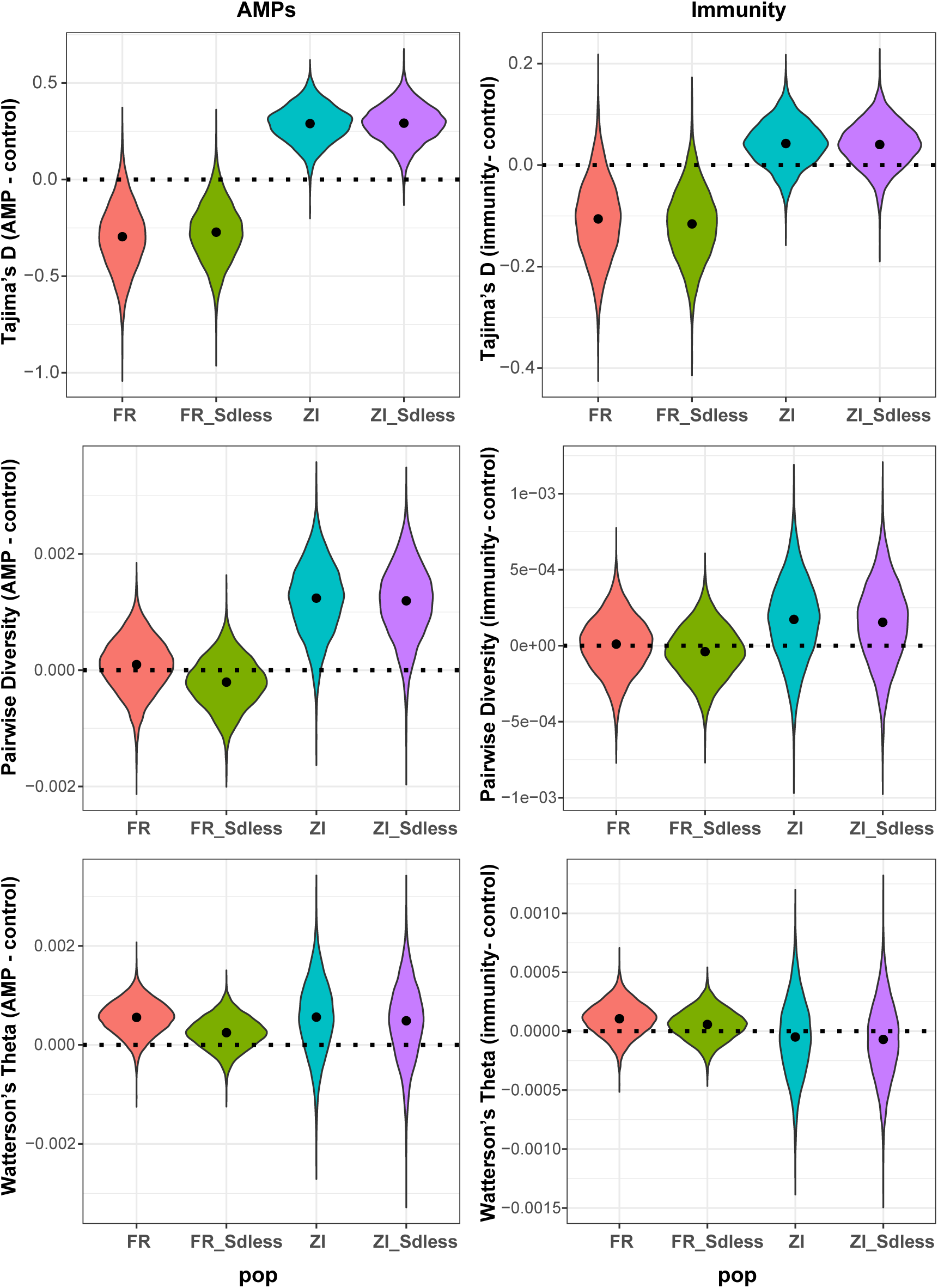
Summary of resampling results (case - control) for AMPs and other immunity genes for Tajima’s D, *π*, Watterson’s *θ*, comparing the results of ZI and FR populations with and without SD chromosomes.

**Supplementary Figure 4.**
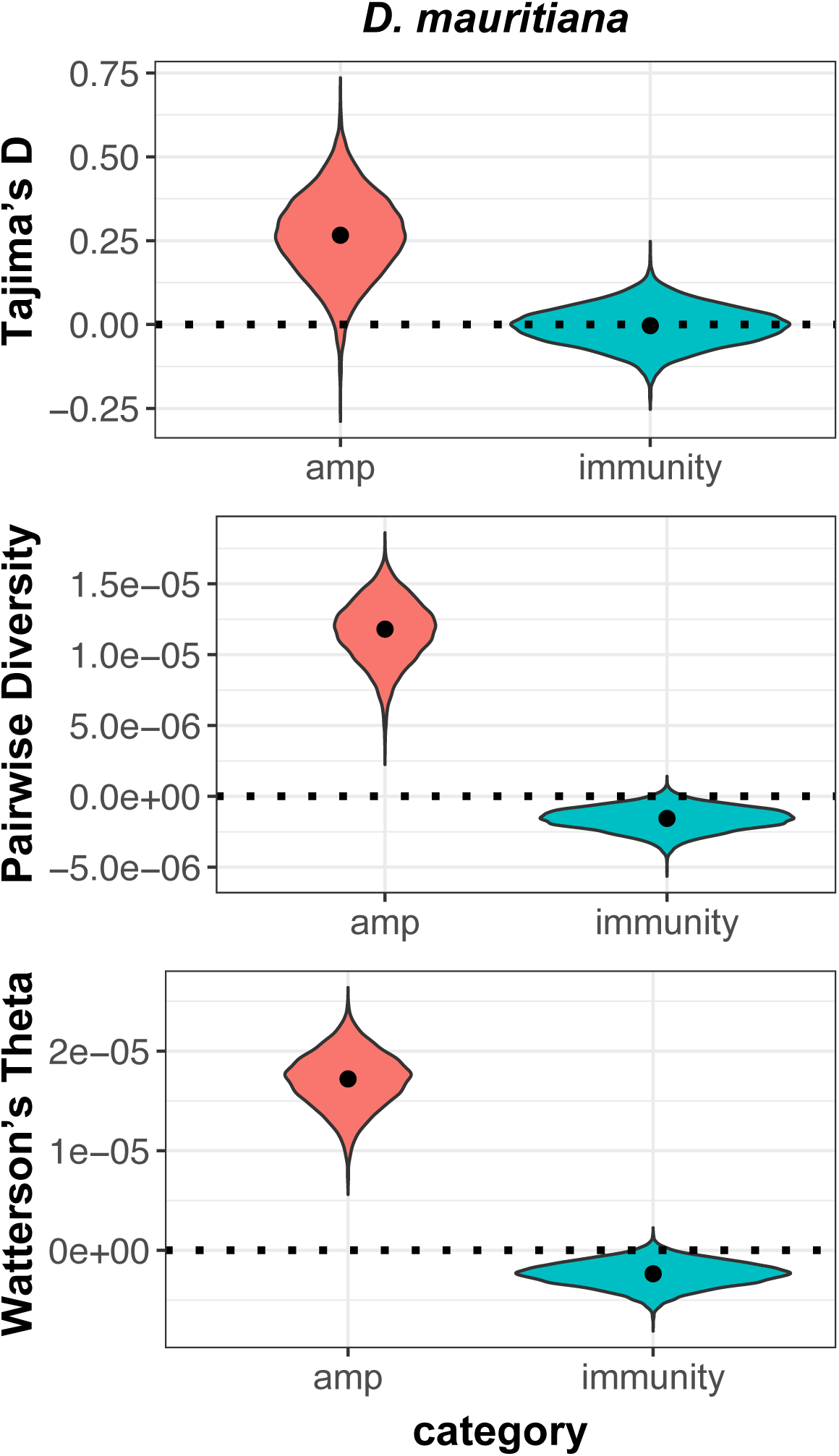
Summary of resampling results (case - control) for AMPs and other immunity genes for Tajima’s D, *π*, Watterson’s *θ* in the *D. mauritiana* population.

## 6 Acknowledgments

We would like to thank John Kelly and Jamie Walters for thoughtful feedback on a previous version of the manuscript. Josep Comeron graciously provided additional data files from his background selection paper. Michael Degiorgio and Katie Siewart provided guidance on using their software packages for *Drosophila* populations. This work was supported by the National Institutes of Health (P20 GM103420 and R00 GM114714 to RLU) and the Max Kade Foundation (fellowship to TH). The funders had no role in study design, data collection and interpretation, or the decision to submit the work for publication.

